# Effects of Transcranial Direct Current Stimulation on GABA and Glutamate in Children: A Pilot Study

**DOI:** 10.1101/759290

**Authors:** C Nwaroh, A Giuffre, L Cole, T Bell, H. L. Carlson, F. P. MacMaster, A Kirton, AD Harris

## Abstract

Transcranial direct current stimulation (tDCS) is a form of non-invasive brain stimulation that safely modulates brain excitability and has therapeutic potential for many conditions. Several studies have shown that anodal tDCS of the primary motor cortex (M1) facilitates motor learning and plasticity, but there is little information about the underlying mechanisms. Using magnetic resonance spectroscopy (MRS) it has been shown that tDCS can affect local levels of γ-aminobutyric acid (GABA) and Glx (a measure of glutamate and glutamine combined) in adults, both of which are known to be associated with skill acquisition and plasticity; however this has yet to be studied in children and adolescents. This study examined GABA and Glx in response to conventional anodal tDCS (a-tDCS) and high definition tDCS (HD-tDCS) targeting the M1 in a pediatric population. Twenty-four typically developing, right handed children ages 12–18 years participated in five consecutive days of tDCS intervention (sham, a-tDCS or HD-tDCS) targeting the right M1 while training in a fine motor task (Purdue Pegboard Task) with their left hand. Glutamate and GABA were measured before and after the protocol (at day 5 and 6 weeks) using conventional MRS and GABA-edited MRS in the sensorimotor cortices. Glutamate measured in the left sensorimotor cortex was higher in the HD-tDCS group compared to a-tDCS and sham at 6 weeks (p = 0.001). No changes in GABA were observed in either sensorimotor cortex at any time. These results suggest that neither a-tDCS or HD-tDCS locally affect GABA and glutamate in the developing brain and therefore it may demonstrate different responses in adults.

## Introduction

Transcranial direct current stimulation (tDCS) is a form of non-invasive brain stimulation in which a weak electrical current is passed between two electrodes placed on the scalp. Using various tDCS montages, cortical excitability can shift to a state of excitation (anodal tDCS) or inhibitory (cathodal tDCS). Placing the anode electrode over M1 for instance typically increases cortical excitability in M1 (1–3). Previous research suggests that changes in excitability outlasts the stimulation session by up to 90 minutes (2, 4). The prolonged and promising changes in both cortical excitability and promising changes in behavioral outcomes combined with its simple application and low cost makes tDCS an attractive as a possible therapeutic tool for a range of clinical conditions (5). For example, tDCS has been suggested to improve symptoms and/or assist in rehabilitation for many neurological disorders with minimal side effects (6), including migraine (7), stroke (8), Parkinson’s disease (9), pain disorders (10) and neurodegenerative disorders (11), as well as psychiatric disorders including depression (12).

High definition tDCS (HD-tDCS) is a newer, more focal form in tDCS that uses arrays of smaller electrodes to improve stimulation localization (13). Most typically used is the 4 x 1 configuration where a central electrode, which determines montage polarity, is placed over the target cortical region, and four outer electrodes (arranged as a ring), act as the reference electrodes. The radii of the surrounding reference electrodes define the region undergoing modulation (14). This configuration has been shown to modulate excitability in a smaller, more specific region compared to conventional tDCS (14, 15). In addition to a more focussed current, its effects on patterns of cortical excitability in the M1 outlast those induced by conventional tDCS, as quantified by motor evoked potentials in response to stimulation (16). Studies support its tolerability in both healthy subjects and patients at intensities up to 2 mA for up to 20 minutes (15–17).

Few studies have investigated tDCS in children, despite its potential (18–21). tDCS administered in a multiday paradigm to the M1 of healthy children while performing a motor task demonstrated greater increases in motor skill compared to sham and improvements are retained 6 weeks later (22, 23). These findings suggest the potential utility of tDCS as a therapeutic tool in children with motor impairments but the biological mechanisms behind these effects remain unknown (24).

Adult studies using magnetic resonance spectroscopy (MRS) to measure regional brain metabolites typically show a decrease in GABA (4, 25, 26) and an increase in Glx (glutamate and glutamine in combination) (4, 26, 27) in the sensorimotor cortex following M1 anodal stimulation. Both GABA, a major inhibitory neurotransmitter, and glutamate, a major excitatory neurotransmitter, are mediators in long-term potentiation (28, 29) and have been associated with behavioral changes following anodal tDCS, quantified as changes in task performance (4, 25, 30). However, it is unknown if these finding translate to a pediatric population and how long these changes in metabolites persist.

Conventional MRS at 3T measures glutamate, though it is often reported as Glx, representing the combination of glutamate and glutamine as their spectra are highly overlapped, making it difficult to reliably resolve these two signals. GABA, on the other hand, is at low concentration and its signal is overlapped by more abundant metabolites and therefore requires editing for accurate measurement (31). GABA-edited MEGA-PRESS, selectively manipulates the GABA signal at 3 ppm by applying an editing pulse to the coupled GABA signal at 1.9 ppm in half of the averages (ON), which are interleaved with averages in which the editing pulse is applied elsewhere not coupled to GABA (OFF). The difference spectrum is acquired by subtracting the ON from the OFF, which removes all peaks not affected by the 1.9 ppm editing pulse (specifically the 3 ppm creatine peak), revealing the GABA signal at 3 ppm.

In this study, GABA-edited and conventional MRS were used to investigate changes in GABA and Glx in response to anodal tDCS (a-tDCS) and anodal HD-tDCS in a pediatric population. By observing metabolite changes in the targeted right sensorimotor cortex and the contralateral left sensorimotor cortex, we aimed to gain insight into the metabolite changes induced by tDCS both after stimulation has concluded and at 6 weeks follow up, with the overall goal of better understanding the mechanism by which tDCS modulates motor learning in the developing brain. Based on the adult literature, we expected GABA to decrease following tDCS and at 6-weeks follow up we expect metabolites to return towards baseline with similar results observed for both anodal and high definition tDCS groups.

## Materials and Methods

This study was a component of the Accelerated Motor Learning in Pediatrics (AMPED) study, a randomized, double-blind, single-center, sham-controlled intervention trial registered at clinicaltrials.gov (NCT03193580) with ethics approval from the University of Calgary Research Ethics Board (REB16-2474). Upon enrolment, participants and guardians provided written, informed consent or assent and were screened to ensure they met safety criteria for non-invasive brain stimulation and MRI scanning. Participants were blinded to the experimental group to which they were assigned and only the investigator administering stimulation was aware of the group until all data was collected. Group assignment was only revealed for data analysis after the study was completed. Additional details regarding the parent study design, recruitment and primary motor learning outcomes can be found in Cole and Giuffre et al (23).

### Experimental Design

Twenty-four typically developing right-handed participants ages 12 to 18 were recruited through the Healthy Infants and Children Clinical Research (HICCUP) Database. The Edinburgh Handedness Inventory was used to confirm right hand dominance with a laterality index ≥ −28. Participants were excluded for MRI contraindications, neuropsychiatric or developmental disorder diagnoses, medications or pregnancy. All participants received a baseline MR scan and motor assessments prior to tDCS. Participants were then computer randomized to a single tDCS condition (n = 8 for each intervention group) with the anode targeting right M1: a-tDCS (1mA conventional anodal tDCS), HD-tDCS (1mA high definition anodal tDCS) and sham tDCS. Participants took part in a 5-day protocol in which they received stimulation each day whilst training in the Purdue Pegboard Task (PPT) using their non-dominant left hand. After stimulation had concluded on Day 5, they received a post-simulation MR scan and completed all motor assessments. Participants returned 6 weeks (± 1 week) later for a follow-up MR scan and motor assessments. The experimental design for this study is shown in Fig 1a.

**Fig 1.**
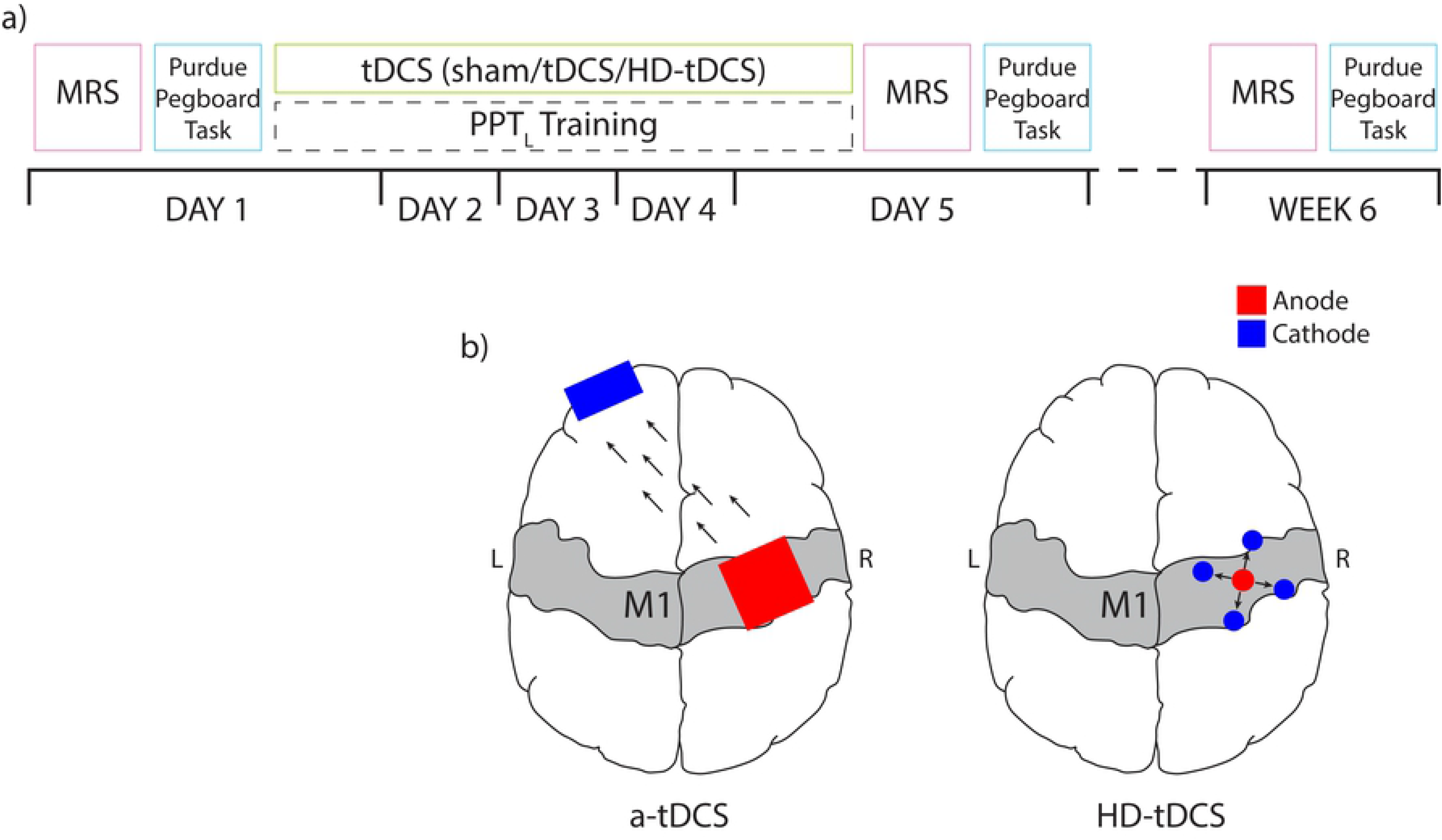
Layout of experimental procedure. a) On Day 1, spectroscopy measurements were collected followed by the Purdue Pegboard Task. Participants then underwent five consecutive days of right M1 targeted anodal tDCS paired with left hand motor training. Participants repeated Day 1 assessments after intervention on Day 5 and at 6 week follow up. b) Anodal tDCS electrode montages shown for a-tDCS (left) and HD-tDCS (right) intervention groups where the anode is red, the cathode is blue and current flow is illustrated with black arrows. MRS, magnetic resonance spectroscopy; PPT, Purdue pegboard task, tDCS, transcranial direct current stimulation; HD-tDCS, high definition tDCS.

### Transcranial Direct Current Stimulation

Participants received 20 minutes of 1mA anodal tDCS in a montage dependent on the assigned stimulation condition. tDCS was administered using a conventional 1 x 1 tDCS or a 4 x 1 HD-tDCS system (Soterix Medical Inc., New York, USA) (Fig 1b). For participants in the a-tDCS or sham group, two 25 cm^2^ saline-soaked sponge electrodes were held on the scalp using a light plastic headband (SNAPstrap, Soterix Medical Inc., New York, USA). The active (anodal) electrode was centred on the right M1 (identified using robotic single pulse transcranial magnetic stimulation;TMS) and the cathodal electrode was placed on the contralateral supraorbital notch. The electrodes were connected to a 1 x 1 DC SMARTscan Stimulator (Soterix). This montage demonstrated in Fig 1b has been used extensively in tDCS studies for motor training in the nondominant left hand (4, 22, 23, 25, 32, 33).

For the HD-tDCS group, a 10:20 EEG cap was used to center the anodal electrode on the right M1, after identifying the location with single pulse TMS as above. The four cathodes were placed ~5 cm away in a 4 x 1 configuration (Fig 1b) using a 4 x 1 HD-tDCS Adaptor and a SMARTscan Stimulator (Soterix) as described previously (15, 34, 35).

For the active stimulation conditions (a-tDCS and HD-tDCS), current was ramped up to 1 mA over 30 seconds and remained at 1mA for 20 minutes. The current was then ramped back down to 0 mA over 30 seconds. For the sham stimulation condition, current was ramped up to 1 mA over 30 seconds and then immediately ramped back down to 0 mA over 30 seconds. After 20 minutes, current was ramped up to 1 mA and then back down to 0 mA over 30 seconds. This procedure is used to mimic the sensations associated with active stimulation and has been previously validated (36). During the 20 mins of stimulation (or sham) participants performed the Purdue Pegboard Task with their left hand (PPT_L_) three every 5 minutes.

### Motor Assessments

The motor assessment was the Purdue Pegboard Task (PPT) (37). This test uses a rectangular board with two sets of 25 holes running vertically down the board and four concave cups at the top of the board that contain small metal pegs. Subjects are asked to remove pegs form the cups and place them in the holes one-at-a-time, as quickly as possible. This task challenges hand dexterity and coordination. A score is given as the number of pegs successfully placed in the holes in 30 seconds with the left hand (PPT_L_). Secondary assessments were the performance of this task with the right hand (PPT_R_) or bimanually (PPT_LR_). Changes in score is reported as ΔPPT.

### MRS Acquisition

Spectroscopy data was collected before the tDCS intervention (baseline), after 5 days of tDCS paired with motor training, and at 6-weeks after tDCS in all 24 subjects on a 3T GE MRI scanner equipped with a 32-channel head coil. Axial T1-weighted fast spoiled gradient recalled echo (FSPGR) brain volume images (BRAVO) were acquired (TR = 7.4 ms, TE = 2.8 ms with 1 mm^3^ voxels) for voxel placement and tissue segmentation. Metabolites were measured in 30 × 30 × 30 mm^3^ voxels located on the right and left sensorimotor cortices. The sensorimotor cortex was identified by Yousry’s hand-knob (38) and the voxel was rotated to align with the cortical surface (Fig 2). GABA data were acquired using a MEGA-PRESS sequence with the following parameters: TR/TE=1800/68 ms, 256 averages; 14 ms editing pulses applied at 1.9 ppm and 7.46 ppm alternating every two averages, and 16 unsuppressed water scans. A conventional PRESS sequence was used to acquire MRS data from which glutamate (as Glx) was quantified with the following parameters: TR/TE=1800/35 ms, 64 averages and 8 unsuppressed water scans. In order to perform symmetrical assessment of the left and right sensorimotor cortices, the water-fat shift directions were mirrored for the sensorimotor voxels for both the GABA-edited MRS and the PRESS acquisitions.

**Fig 2.**
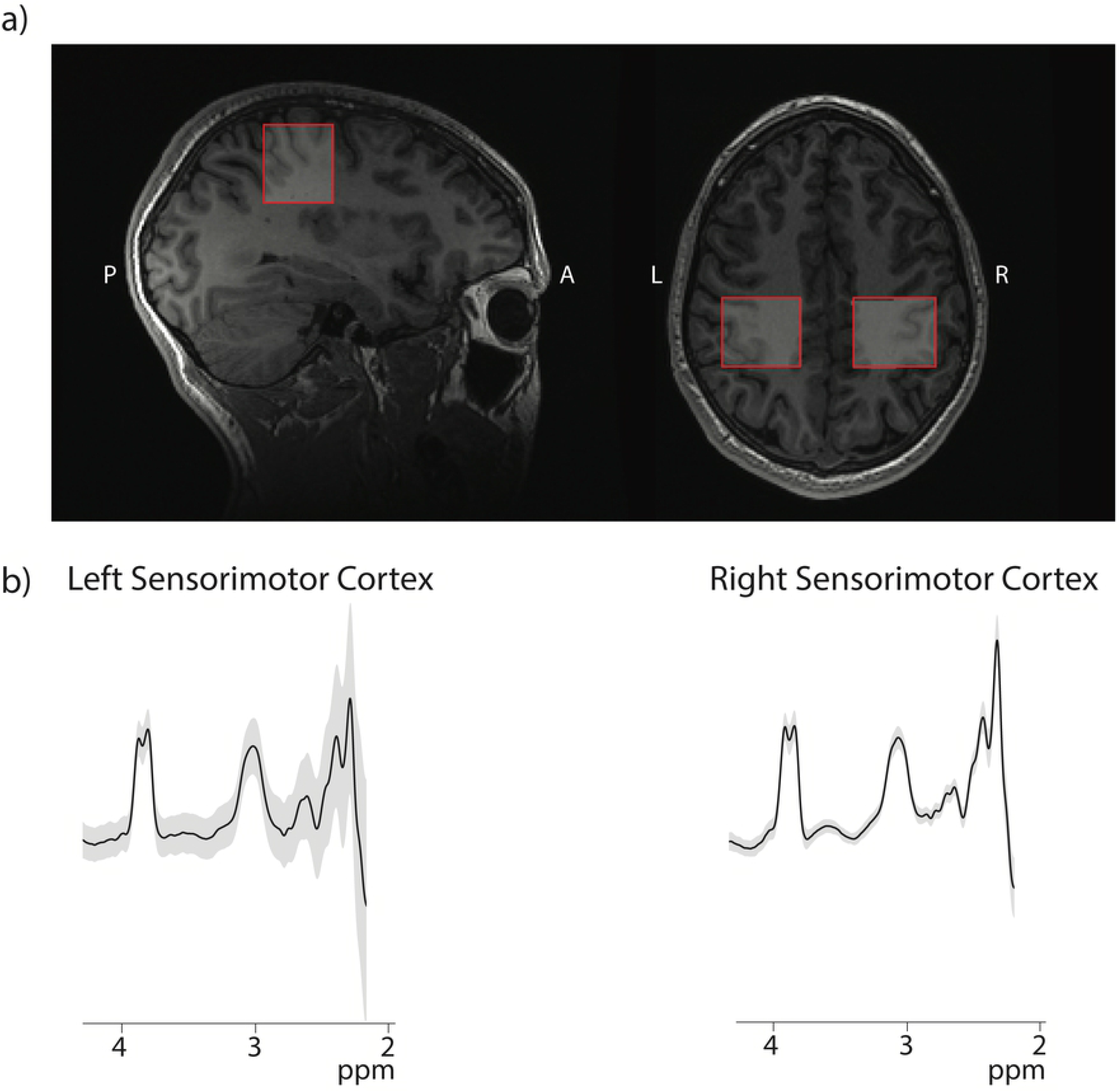
Voxel Placement and Data Quality. Example of voxel placement in the sensorimotor cortex on a participant T1-weighted image. b) MEGA-PRESS spectra acquired in each location. The black line depicts the average fit line and the grey area shows ±1 standard deviation in the right and left sensorimotor cortex.

### MRS data analysis

GABA data were analyzed using GANNET 3.0 (39) software in MATLAB R2014a (The Mathworks, Natick, MA, USA), including retrospective frequency and phase correction and correction for voxel tissue content, assuming grey matter contains twice as much GABA as white matter (i.e., α = 0.5 as per literature) (40). In this experiment, we assumed sensorimotor voxels were composed of 40% grey matter and 60% white matter in the GABA tissue correction (41). Conventional PRESS data was corrected for frequency and phase drift using the FID-A toolkit (42) and then analyzed using LCModel (43) with basis sets developed from LCModel. Metabolite levels from LCModel were tissue-corrected using the Gasparovic approach (44) and the CSF voxel fraction, accounting for the negligible metabolites present in CSF. As a confirmatory analysis, metabolite levels referenced to creatine were also examined.

### Statistical Analysis

All statistical analyses were performed using SPSS Statistics 25 (IBM, Armonk, NY, USA). Demographic data of the three groups (a-tDCS, HD-tDCS and sham) were compared with an ANOVA model and Chi-squared for sex data. Changes in GABA and glutamate between tDCS conditions and over time were assessed using a linear mixed model analysis with fixed effects for intervention and experimental day, the interaction of intervention and experimental day, and covariates for age and sex for each voxel. Post-hoc pair-wise analyses with Bonferroni correction for multiple comparisons were performed to specifically examine effects of intervention and experimental day.

Partial correlations controlling for intervention were used to examine the relationship between changes in metabolites and changes in motor assessment performance before and after stimulation, and 6 weeks after stimulation had concluded. Initially these correlations were pooled across all groups and follow-up analyses were performed in each group as appropriate.

## Results

### Population Characteristics

Twenty-four typically developing children (mean 15.5 ± 1.7 years, 13 females and 11 males) completed all phases of the study with no drop outs. Due to technical difficulties, one participant did not have GABA or Glx data available in both sensorimotor cortices in the post intervention timepoint. Population demographics are shown in Table 1. Age, sex and laterality index did not differ significantly between groups (p > 0.3 for all parameters).

**Table 1.** Mean participant demographics ± 1 standard deviation for all stimulation intervention groups. No significant difference between groups was identified.

| | **SHAM**<br>(± SD) | **A-TDCS**<br>(± SD) | **HD-TDCS**<br>(± SD) | **MEAN**<br>(± SD) | **BETWEEN**<br>**GROUPS** |
| --- | --- | --- | --- | --- | --- |
| **AGE** | 15.81<br>(± 1.3) | 15.94<br>(± 1.5) | 14.77<br>(± 2.0) | 15.51<br>(± 1.7) | p = 0.324 |
| **LATERALITY**<br>**INDEX** | 81.9<br>(± 22.8) | 82.5<br>(± 13.1) | 81.3<br>(± 14.7) | 81.9<br>(± 16.6) | p = 0.879 |
| **SEX (F:M)** | 6:2 | 3:5 | 4:4 | 13:11 | p = 0.309 |

### Data Quality

The GABA-edited spectra from the right and left sensorimotor cortices from all time points are show in Fig 2b; the grey shows a single standard deviation range across all data and the black line is the average of all data. All data, both GABA-edited and conventional PRESS, were assessed for quality by visual inspection as well as a CRLB threshold of 20%. One PRESS dataset was excluded due to poor data quality, the remaining spectra were of high quality with a mean SNR of 41.4 ± 6.3, all FWHM water <15 Hz, mean FWHM water 6.01 ± 1.92 Hz. MEGA-PRESS GABA data was also of high quality across all data sets: all fit errors < 10%, mean fit error 4.59 ± 1.21, all FWHM Cr <10%, mean FWHM Cr: 9.57 ± 0.92 Hz. Generally, spectra with fit errors below 12% are deemed to be of sufficient quality (39).

### Glx and GABA Group Changes

Linear mixed model analyses showed a significant fixed effect of tDCS intervention over time on Glx levels in the left sensorimotor cortex (p = 0.010). Post-hoc Bonferroni corrected pairwise analyses showed at the 6 week follow up, Glx was significantly higher in the HD-tDCS group compared to the sham group (p = 0.001; Fig 3). In the HD-tDCS group, Glx in the left sensorimotor cortex increased between post-intervention and the 6 week follow up time points (p = 0.042), however, this did not withstand correction for multiple comparisons (Fig 3). No significant fixed effect of tDCS intervention over time for Glx was detected in the right sensorimotor cortex (p = 0.221). No significant fixed effect was observed in either sensorimotor cortex (left: p = 0.248; right: p = 0.724) for GABA. Metabolite data referenced to creatine showed the same results. No significant metabolite change difference were detected between that a-tDCS and sham groups in both left and right sensorimotor cortices.

**Fig 3.**
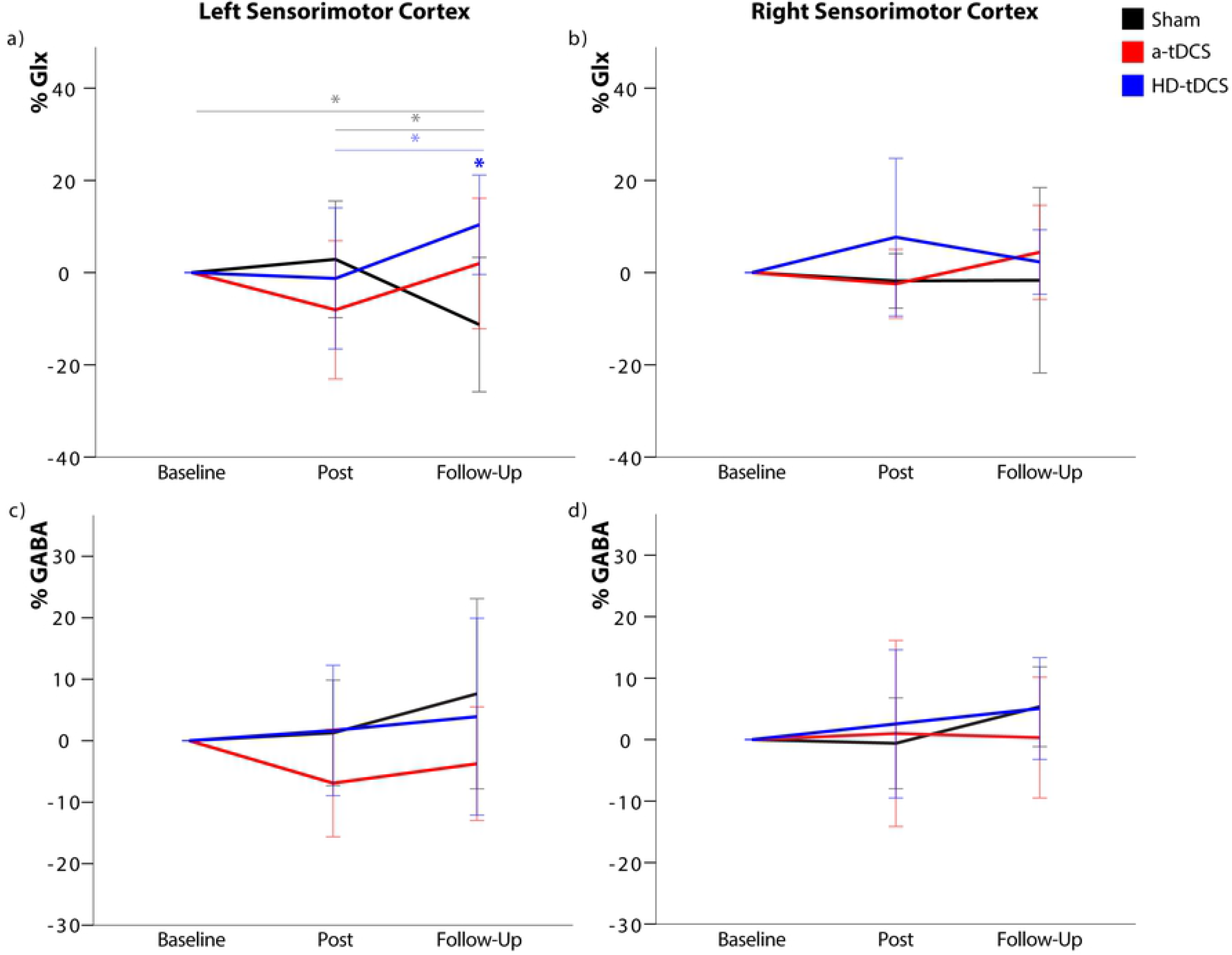
Metabolite Changes Over Time. Changes in metabolite levels for all intervention groups (sham in black, tDCS in red and HD-tDCS in blue) over the duration of the experiment given as a percentage change from baseline values (mean ± 1 SD). * p > 0.05, those in bold withstand Bonferroni correction for multiple comparisons while those that are transparent lose significance following multiple comparisons correction.

### Relationship between metabolite changes and motor performance

Partial correlation analysis comparing changes in GABA and Glx, pooled across the three intervention groups, showed a significant positive relationship between the change in left sensorimotor GABA (%GABA) and change in PPT_L_ score (ΔPPT_L_) (r = 0.538, p = 0.018; Fig 4d), participants with a greater positive change in GABA showed a greater improvement in PPT. Post-hoc assessments by intervention groups showed this relationship was maintained in the anodal tDCS group only (r = 0.864, p = 0.006; Fig 4d).

**Fig 4.**
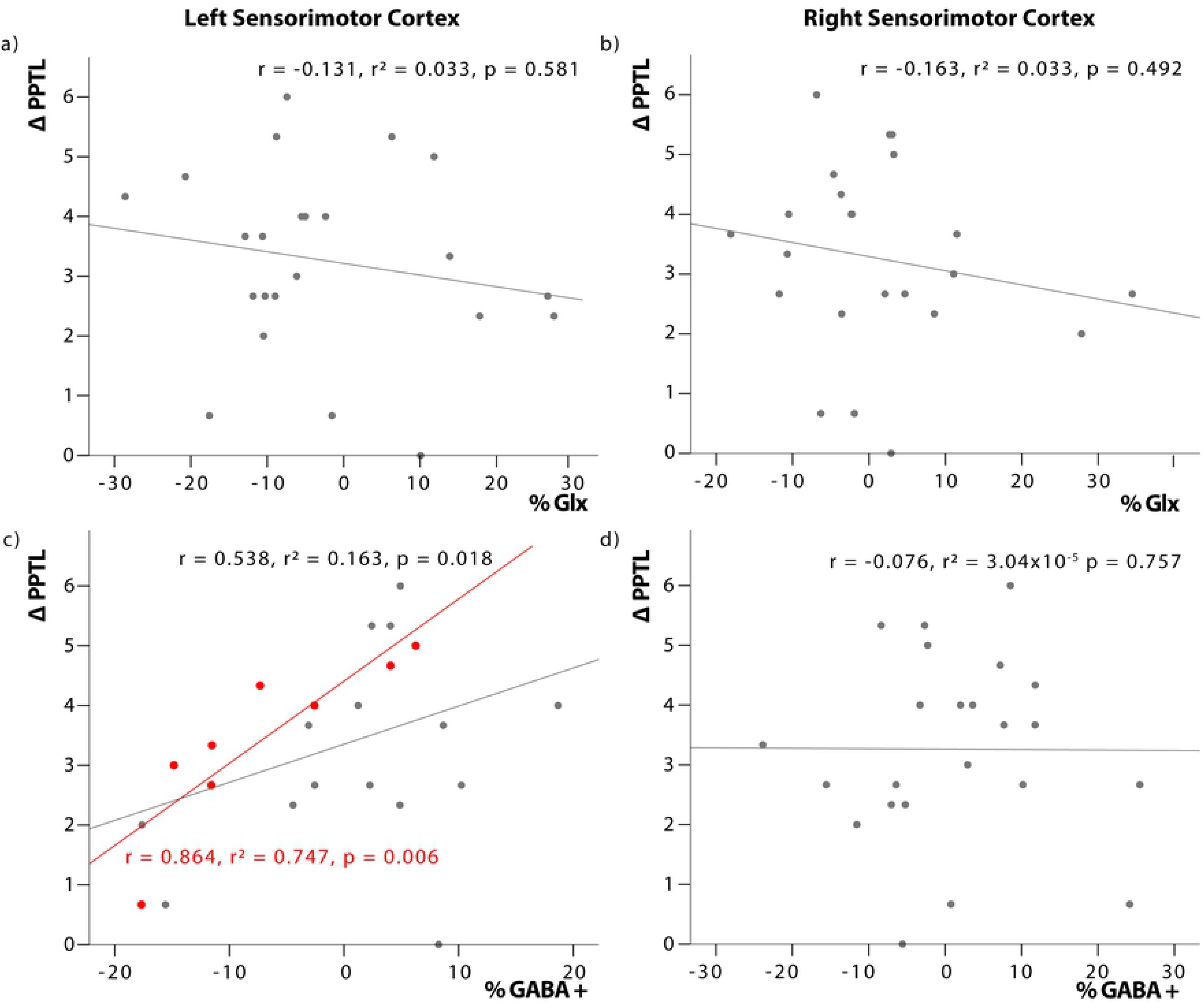
Relationship between changes in metabolite concentration and motor performance. Correlationn between change in metabolite concentration (% Glx and %GABA) and change in Purdue Pegboard Task post intervention (ΔPPTL) controlling for intervention group and age. Left sensorimotor cortex GABA is significantly correlated with PPTL for the pooled intervention groups (grey line). This relationship is also observed in the anodal tDCS intervention group (red).

No significant relationship was observed between ΔPPT_L_ and changes in GABA in the right sensorimotor cortex (r = −0.065, p = 0.784; Fig 4c). Additionally, no significant relationship was seen between changes in PPT score and changes in Glx in the right (Fig 4a) or left (Fig 4b) sensorimotor cortex (p > 0.05).

## Discussion

Several adult studies have shown that single (43, 44) or multiple session (30, 45) tDCS paired with training in a motor task is associated with improvements in said task and improvements in performance are greater than motor training alone (i.e., sham-tDCS). The same is observed in pediatric studies (22, 23), however results may differ slightly in terms of the phase of learning affected by stimulation. Results in children suggest that tDCS facilitates online learning (22) while in adults evidence suggests tDCS enhances learning primarily through offline effects (30). GABA and glutamate are involved in learning (24, 28, 46) and have both been observed to change in response to anodal tDCS in adults (4, 24–26, 46, 47). This study examined changes in GABA and Glx in response to right M1 anodal tDCS and HD-tDCS in a pediatric population. Metabolites were measured at baseline, after a 5-day tDCS and motor learning intervention (post-intervention) and at 6 weeks follow-up.

To our knowledge, this is the first investigation of metabolite changes in response to tDCS in a typically developing pediatric population. Additionally, this is the first-time metabolites have been measured in a control population after a multiday protocol with a follow-up assessment. Previous studies in adults have illustrated that GABA decreases (33, 46) and glutamate increases (47), with skill acquisition and improved function in the region responsible for the skill execution, the M1. It has been suggested that tDCS facilitates changes in GABA and glutamate to augment learning. Studies conducted in adults have shown anodal tDCS increases sensorimotor glutamate (4, 26, 27) and decreases GABA (4, 25, 26, 48); however, others have failed to replicate these findings. Similarly, we did not see decreased GABA and increased Glx at the site of stimulation, though we did see contralateral changes. Our results potentially indicate the developing brain responds differently to tDCS compared to the adult brain.

### Post-Intervention Changes in GABA and Glx

Following five days of tDCS and motor training there were no significant changes in metabolite levels in either the right or left sensorimotor cortex, though trends toward decreased left sensorimotor GABA (contralateral to the tDCS target) in the a-tDCS group were seen. Adult literature using healthy controls suggests acute decrease in GABA local to the tDCS target (4, 25, 26, 48). Similarly, participants with a neurodegenerative condition who followed a protocol of 15 a-tDCS sessions also showed decreased GABA in the tissue targeted with a-tDCS (11). Given the contrast of our results and those in the literature, we suggest that the pediatric brain responds differently to tDCS.

In healthy adults, GABA and glutamate in the motor cortex work together to maintain an excitation-inhibition balance that is crucial for plasticity (49). It has been suggested that this balance of GABA and glutamate can be shifted to a relative optimum level that is thought to mediate behavioral outcomes (50). It is possible that in the developing brain, this excitation/inhibition balance is more dynamic while in the adult brain it is relatively static. When an external stimulus is introduced, like tDCS or a foreign motor task, the adult brain shows a shift to facilitate plasticity while the pediatric brain was already in its “plastic state”. There is also evidence describing the pediatric brain as being hyperexcitable (19) which may suggest it has a lower concentration of GABA (51, 52), and therefore less dynamic range to reduce GABA compared to the adult brain where increased GABAergic inhibition is necessary to refine already acquired skills.

Secondly, transcallosal inhibitory processes (53) may have a more pronounced effect in the pediatric brain. Here we show trends towards decreased GABA in the left sensorimotor cortex, contralateral to the site of stimulation, as opposed to changes in the site of stimulation (right cortex). This suggests lateralization of motor learning in the left dominant cortex as previously described by Schambra et al (54). The impact of transcallosal inhibition is also seen in pediatric studies applying tDCS contralateral to stroke lesions in an effort to augment motor learning of the affected hemisphere (55, 56). According to pediatric models of anodal tDCS, the current appears to travel through the motor fibers of the corpus callosum into the contralateral hemisphere (56). However, the same mechanism is not expected to be true for HD-tDCS which has a more focal current.

Finally, as mentioned above, tDCS may act on different phases of learning in children compared to adults, therefore the paradigm in which we expect GABA and glutamate changes to appear shortly after stimulation is not the appropriate time window to detect changes. Similarly, it is possible that the metabolic response to stimulation changes with applications over consecutive days. In this study, we suspect participants may have transitioned into a phase of learning that requires less plasticity and the cortex is no longer responding to tDCS with the predicted GABA and Glx changes at five days when our measures were taken. Adult literature suggests the changes in GABA and glutamate measured by MRS in response to learning vary with time (46, 57) and it is possible that a ceiling of PPT skill, and also of metabolite change, was reached before our MRS measurements were taken.

Although no significant changes in GABA concentration were detected between MRS measurements, this does not conclusively rule out GABAergic changes in response to motor learning. It is possible that subtle biphasic changes in GABA are taking place during motor learning that we are unable to detect. While this cannot be confirmed in our investigation, there is literature suggesting changes in GABA concentration are time sensitive with fluctuation in GABA concentration occurring in the 90 minute window following stimulation (4, 46, 57). The time sensitivity of metabolite measurements is further supported by seemingly discrepant findings in the literature in which GABA and Glx changes are not seen during tDCS (58–60).

Changes in Glx in response to stimulation in the literature are inconsistent. Clark et al. reports Glx increases after anodal tDCS and suggest that tDCS may involve the NMDA pathway (27). Stagg et al. also reports changes in Glx in response to cathodal tDCS (4). They propose MRS measures of Glx lack sensitivity to consistently detect Glx changes following tDCS (4, 25). Several other studies report an absence of significant changes in Glx in response to a-tDCS with little speculation as to why (4, 26, 58, 59, 61).

### 6 Week Follow Up in GABA and Glx

At 6 weeks follow up, it was expected that metabolites would return to baseline to maintain homeostatic balance in the brain after the initial phases of skill acquisition had concluded, while retaining motor skill improvements. However, we observed a significant increase in the left sensorimotor Glx at 6 weeks follow up in the HD-tDCS group compared to the sham group (p = 0.001) and compared to HD-tDCS baseline level. We also see a trend of increased Glx in the HD-tDCS group between post intervention and 6-week follow up in the left sensorimotor cortex (23). This suggests that in the hemisphere contralateral to stimulation, HD-tDCS has a longer-term modulation of glutamatergic pathways. When examined in conjunction with the secondary motor data collected, the change in left sensorimotor Glx in the HD-tDCS group is accompanied by an improvement in the right hand PPT at 6 weeks follow up. These results can be explained by the motor overflow mechanism. Motor overflow is a phenomena that typically disappears in late childhood and describes unintentional movement that mirror voluntary movements typically in homologous muscle on the opposite side of the body (62). Similarly, the decrease in GABA in the left sensorimotor cortex in the a-tDCS group persisted. Persistent decreases in GABA several weeks after tDCS intervention have been seen in primary progressive aphasia (11); though those changes were seen at the site of stimulation. Several studies have shown improvement in motor learning in the contralateral hand following tDCS of either the right or left M1 (63–65). The “callosal access” hypothesis dictates that performance can be facilitated in the untrained limb due to motor engrams developed in the dominant hemisphere. These engrams underlie performance of the trained hand located in homologous regions that the opposite motor cortex can access via the corpus callosum (55, 66, 67).

### Relationship Between Changes in Metabolites and Changes in Motor Performance

We found a significant, positive relationship between change in left sensorimotor GABA (cortex contralateral to stimulation) and improvement in the task performance by the left hand post tDCS intervention and training, further supporting the above mentioned callosal hypothesis. Those participants who experience a greater positive change in GABA concentration in the hemisphere contralateral to stimulation (left motor cortex) present a greater improvement in PPT score over the 5-day stimulation and training period. This relationship is specifically seen in the a-tDCS group only, suggesting that anodal stimulation induces a contralateral inhibition that does not occur with HD-tDCS or in normal (sham group) learning, driving an enhanced improvement in PPT score.

No relationship between changes in Glx and task performance post-intervention nor between GABA or Glx and change in PPT score 6 weeks after stimulation and training was observed. These results are in accordance with adult studies that report no significant relationship between change in motor skill and concentration of Glx in the motor cortex contralateral to the hand executing the task (33). However, adult studies have reported a relationship between task improvement and GABA changes in the tDCS targeted cortex (i.e. right sensorimotor GABA changes and left hand training and task performance) (25, 33). This dissimilarity suggests that neurochemistry in the pediatric and adult brain respond in different ways during motor learning, warranting further investigation.

## Conclusions

Non-invasive stimulation is an expanding area of research with investigations into the use of modalities similar to tDCS being investigated as a therapy for a range of disorders including migraine, pain and stroke (6, 7, 9, 11, 12, 18, 67). While these studies have suggested that non-invasive brain stimulation can improve outcomes, there is little analysis into the underlying physiological changes behind these responses are not well understood, particularly in the developing brain. This study aimed to shed light on the metabolite changes induced by M1 anodal tDCS in conjunction with a motor training paradigm.

We investigated changes, in GABA and glutamate concentrations following 5 consecutive days tDCS comparing conventional anodal tDCS, HD-tDCS and sham. Unexpectedly, Transcranial direct current stimulation (tDCS) produces localized and specific alterations in neurochemistry: A 1H magnetic resonance spectroscopy study significant changes in metabolites at the site of stimulation post 5-day tDCS intervention or 6 weeks after the intervention. It is possible that changes in metabolites occur immediately after stimulation and learning and this effect is diminished over the 5 days stimulation as skill level improves. However, we suggest the pediatric brain responds differently to tDCS compared to adults. In particular, we suggest contralateral modulation of learning and metabolites has a greater role in the pediatric brain, highlighting the need for further study of the effects of non-invasive stimulation on the pediatric brain specifically. Furthermore, we also show the response to HD-tDCS is different compared to a-tDCS based on the observation of increased glutamate in the left sensorimotor cortex 6 weeks after stimulation specifically in response to HD-tDCS. Further investigation into the effects of HD-tDCS is needed to determine its efficacy on motor learning.

## References

1. Reis J, Fritsch B. Modulation of motor performance and motor learning by transcranial direct current stimulation. Curr Opin Neurol. 2011;24(6):590–6.

2. Nitsche MA, Paulus WJ. Sustained excitability elevations induced by transcranial DC motor cortex stimulation in humans. Neurology. 2001;57(10):1899–901.

3. Nitsche MA, Schauenburg A, Lang N, Liebetanz D, Exner C, Paulus WJ, et al. Facilitation of implicit motor learning by weak transcranial direct current stimulation of the primary motor cortex in the human. J Cogn Neurosci. 2003;15(4):619–26.

4. Stagg CJ, Best JG, Stephenson MC, O’Shea J, Wylezinska M, Kincses ZT, et al. Polarity-Sensitive Modulation of Cortical Neurotransmitters by Transcranial Stimulation. J Neurosci. 2009;29(16):5202–6.

5. Zhao H, Qiao L, Fan D, Zhang S, Turel O, Li Y. Modulation of Brain Activity with Noninvasive Transcranial Direct Current Stimulation (tDCS): Clinical Applications and Safety Concerns. 2017;8(May).

6. Nitsche MA, Cohen LG, Wassermann EM, Priori A, Lang N, Antal A, et al. Transcranial direct current stimulation: State of the art 2008. Brain Stimul. 2008;1(3):206–23.

7. Antal A, Lang N, Boros K, Nitsche MA, Siebner HR, Paulus W. Homeostatic metaplasticity of the motor cortex is altered during headache-free intervals in migraine with aura. Cereb Cortex. 2008;18(11):2701–5.

8. Boggio PS, Nunes A, Rigonatti SP, Nitsche MA, Pascual-Leone A, Fregni F. Repeated sessions of noninvasive brain DC stimulation is associated with motor function improvement in stroke patients. Restor Neurol Neurosci. 2007;25(2):123–9.

9. Fregni F, Boggio PS, Santos MC, Lima MC, Vieira AL, Rigonatti SP, et al. Noninvasive cortical stimulation with transcranial direct current stimulation in Parkinson’s disease. Mov Disord. 2006;21(10):1693–702.

10. Fregni F, Boggio PS, Lima MC, Ferreira MJ., Wagner T, Rigonatti SP, et al. A sham-controlled, phase II trial of transcranial direct current stimulation for the treatment of central pain in traumatic spinal cord injury. Pain. 2006;122(1):197–209.

11. Harris AD, Wang Z, Ficek B, Webster K, Edden RA, Tsapkini K. Reductions in GABA following a tDCS-Language Intervention for Primary Progressive Aphasia. Neurobiol Aging [Internet]. 2019;79:75–82. Available from: https://linkinghub.elsevier.com/retrieve/pii/S0197458019300892

12. Fregni F, Boggio PS, Nitsche MA, Marcolin MA, Rigonatti SP, Pascual-Leone A. Treatment of major depression with transcranial direct current stimulation. Bipolar Disord. 2006;8:203–4.

13. Dmochowski JP, Datta A, Bikson M, Su Y, Parra LC. Optimized multi-electrode stimulation increases focality and intensity at target. J Neural Eng. 2011;8(4).

14. Datta A, Bansal V, Diaz J, Patel J, Reato D, Bikson M. Gyri–precise head model of transcranial DC stimulation. NIH Public Access. 2010;2(4):201–7.

15. Villamar MF, Volz MS, Bikson M, Datta A, DaSilva AF, Fregni F. Technique and Considerations in the Use of 4×1 Ring High-definition Transcranial Direct Current Stimulation (HD-tDCS). J Vis Exp. 2013;(77).

16. Kuo HI, Bikson M, Datta A, Minhas P, Paulus W, Kuo MF, et al. Comparing cortical plasticity induced by conventional and high-definition 4 × 1 ring tDCS: A neurophysiological study. Brain Stimul. 2013;6(4):644–8.

17. Borckardt JJ, Bikson M, Frohman H, Reeves ST, Datta A, Bansal V, et al. A pilot study of the tolerability and effects of high-definition transcranial direct current stimulation (HD-tDCS) on pain perception. J Pain [Internet]. 2012;13(2):112–20. Available from: http://dx.doi.org/10.1016/j.jpain.2011.07.001

18. Bikson M, Grossman P, Thomas C, Zannou AL, Jiang J, Adnan T, et al. Safety of Transcranial Direct Current Stimulation: Evidence Based Update 2016. Brain Stimul. 2016;9(5):641–61.

19. Hameed MQ, Dhamne SC, Gersner R, Kaye HL, Oberman LM, Pascual-Leone A, et al. Transcranial Magnetic and Direct Current Stimulation in Children. Curr Neurol Neurosci Rep. 2017;17(2).

20. Kirton A, Ciechanski P, Zewdie E, Andersen J, Nettel-Aguirre A, Carlson H, et al. Transcranial direct current stimulation for children with perinatal stroke and hemiparesis. Neurology. 2016;88(3):259–67.

21. Rajapakse T, Kirton A. Non-invasive brain stimulation in children: Applications and future directions. Transl Neurosci. 2013;4(2):217–33.

22. Ciechanski P, Kirton A. Transcranial Direct-Current Stimulation Can Enhance Motor Learning in Children. Cereb Cortex. 2017;27(5):2758–67.

23. Cole L, Giuffre A, Ciechanski P, Carlson HL, Zewdie E, Kuo H-C, et al. Effects of High-Definition and Conventional Transcranial Direct-Current Stimulation on Motor Learning in Children. 2018;12:1–12.

24. Kirton A. Modeling developmental plasticity after perinatal stroke: Defining central therapeutic targets in cerebral palsy. Vol. 48, Pediatric Neurology. 2013. p. 81–94.

25. Stagg CJ, Bachtiar V, Johansen-Berg H. The role of GABA in human motor learning. Curr Biol. 2011;21(6):480–4.

26. Kim S, Stephenson MC, Morris PG, Jackson SR. TDCS-induced alterations in GABA concentration within primary motor cortex predict motor learning and motor memory: A 7T magnetic resonance spectroscopy study. Neuroimage. 2014;99:237–43.

27. Clark VP, Coffman BA, Trumbo MC, Gasparovic C. Transcranial direct current stimulation (tDCS) produces localized and specific alterations in neurochemistry: A 1H magnetic resonance spectroscopy study. Neurosci Lett. 2011;500(1):67–71.

28. Froc DJ, Chapman A, Trepel C, Racine RJ. Long-Term Depression and Depotentiation in the Sensorimotor Cortex of the Freely Moving Rat. J Neurosci. 2000;20(1):438–45.

29. Lüscher C, Malenka RC. NMDA Receptor-Dependent Long-Term Potentiation and Long-Term Depression (LTP/LTD). 2012;1–15.

30. Carlson HL, Ciechanski P, Harris AD, MacMaster FP, Kirton A. Changes in spectroscopic biomarkers after transcranial direct current stimulation in children with perinatal stroke. Brain Stimul. 2018;11(1):94–103.

31. Harris AD, Saleh MG, Edden R. Edited 1 H magnetic resonance spectroscopy in vivo: Methods and metabolites. Magn Reson Med. 2017;77(4):1377–89.

32. Reis J, Schambra HM, Cohen LG, Buch ER, Fritsch B, Zarahn E, et al. Noninvasive cortical stimulation enhances motor skill acquisition over multiple days through an effect on consolidation. Proc Natl Acad Sci. 2009;106(5):1590–5.

33. Kolasinski J, Hinson EL, Divanbeighi Zand AP, Rizov A, Emir UE, Stagg CJ. The dynamics of cortical GABA in human motor learning. J Physiol. 2019;597(1):271–82.

34. Caparelli-Daquer EM, Zimmermann TJ, Mooshagian E, Parra LC, Rice JK, Datta A, et al. A Pilot Study on Effects of 4×1 High-Definition tDCS on Motor Cortex Excitability. 2017;20(1):48–55.

35. Richardson J, Datta A, Dmochowski JP, Parra LC, Fridriksson J. Feasibility of using high-definition transcranial direct current stimulation (HD-tDCS) to enhance treatment outcomes in persons with aphasia. NeuroRehabilitation. 2015;36(1):115–26.

36. Ambrus GG, Al-Moyed H, Chaieb L, Sarp L, Antal A, Paulus W. Brain Stimulation The fade-in e Short stimulation e Fade out approach to sham tDCS e Reliable at 1 mA for naïve and experienced subjects, but not investigators. Brain Stimul [Internet]. 2012;5(4):499–504. Available from: http://dx.doi.org/10.1016/j.brs.2011.12.001

37. Tiffin J, Asher EJ. The Purdue Pegboard: norms and studies of reliability and validity. J Appl Psychol. 1948;32(3):234–47.

38. Yousry TA, Schmid UD, Alkadhi H, Schmidt D, Peraud A, Buettner A, et al. Localization of the motor hand area to a knob on the precentral gyrus. A new landmark. Brain. 1997;120(1):141–57.

39. Edden R, Puts N, Harris AD, Barker PB, Evans J. Gannet: A batch-processing tool for the quantitative analysis of gamma-aminobutyric acid-edited MR spectroscopy spectra. J Magn Reson Imaging. 2014;40(6):1445–52.

40. Harris AD, Puts N, Edden R. Tissue correction for GABA-edited MRS: Considerations of voxel composition, tissue segmentation, and tissue relaxations. J Magn Reson Imaging. 2015;42(5):1431–40.

41. Harris AD, Gilbert DL, Horn P, Crocetti D, Cecil KM, Edden RA, et al. Anomalous relationship between sensorimotor GABA levels and task-dependent cortical excitability in children with Attention-deficit/hyperactivity disorder. Ann Neurol. 2019;

42. Near J, Edden R, Evans J, Paquin R, Harris AD, Jezzard P. Frequency and phase drift correction of magnetic resonance spectroscopy data by spectral registration in the time domain. Magn Reson Med. 2015;73(1):44–50.

43. Provencher SW. Estimation of metabolite concentrations from localized in vivo proton NMR spectra. Magn Reson Med. 1993;30(6):672–9.

44. Gasparovic C, Song T, Devier D, Bockholt HJ, Caprihan A, Mullins PG, et al. Use of tissue water as a concentration reference for proton spectroscopic imaging. Magn Reson Med. 2006;55(6):1219–26.

45. Fregni F, Boggio PS, Nitsche M, Bermpohl F, Antal A, Feredoes E, et al. Anodal transcranial direct current stimulation of prefrontal cortex enhances working memory. Exp Brain Res. 2005;166(1):23–30.

46. Floyer-Lea A. Rapid Modulation of GABA Concentration in Human Sensorimotor Cortex During Motor Learning. J Neurophysiol. 2006;95(3):1639–44.

47. Cohen-Kadosh K, Krause B, King AJ, Near J, Cohen-Kadosh R. Linking GABA and glutamate levels to cognitive skill acquisition during development. Hum Brain Mapp. 2015;36(11):4334–45.

48. Bachtiar V, Near J, Johansen-Berg H, Stagg CJ. Modulation of GABA and resting state functional connectivity by transcranial direct current stimulation. Elife. 2015;4(September 2015):1–9.

49. Turrigiano GG, Nelson SB. Hebb and homeostasis in neuronal plasticity. Curr Opin Neurobiol. 2000;10(3):358–64.

50. Krause B, Márquez-Ruiz J, Cohen Kadosh R. The effect of transcranial direct current stimulation: a role for cortical excitation/inhibition balance? Front Hum Neurosci. 2013;7(September):1–4.

51. Swann JW, Brady RJ, Martin DL. Postnatal development of GABA-mediated synaptic inhibition in rat hippocampus. Neuroscience. 1989;28(3):551–61.

52. Silverstein FS, Jensen FE. Neonatal Seizures. Volpe’s Neurol Newborn. 2007;6(2):112–20.

53. Ciechanski P, Zewdie E, Kirton A. Developmental profile of motor cortex transcallosal inhibition in children and adolescents. J Neurophysiol. 2017; 118(1):140–8.

54. Schambra HM, Abe M, Luckenbaugh DA, Reis J, Krakauer JW, Cohen LG. Probing for hemispheric specialization for motor skill learning: a transcranial direct current stimulation study. J Neurophysiol [Internet]. 2011;106(2):652–61. Available from: http://www.physiology.org/doi/10.1152/jn.00210.2011

55. Kirton A, deVeber G, Gunraj C, Chen R. Cortical excitability and interhemispheric inhibition after subcortical pediatric stroke: Plastic organization and effects of rTMS. Clin Neurophysiol [Internet]. 2010;121(11):1922–9. Available from: http://dx.doi.org/10.1016/j.clinph.2010.04.021

56. Ciechanski P, Carlson HL, Yu SS, Kirton A. Modeling Transcranial Direct-Current Stimulation-Induced Electric Fields in Children and Adults. Front Hum Neurosci. 2018;12(July):1–14.

57. Patel HJ, Romanzetti S, Pellicano A, Nitsche MA, Reetz K, Binkofski F. Proton Magnetic Resonance Spectroscopy of the motor cortex reveals long term GABA change following anodal Transcranial Direct Current Stimulation. Sci Rep [Internet]. 2019;9(1):2807. Available from: http://www.nature.com/articles/s41598-019-39262-7

58. Hone-Blanchet A, Edden R, Fecteau S. Online Effects of Transcranial Direct Current Stimulation in Real Time on Human Prefrontal and Striatal Metabolites. Biol Psychiatry [Internet]. 2016;80(6):432–8. Available from: http://dx.doi.org/10.1016/j.biopsych.2015.11.008

59. Ryan K, Wawrzyn K, Gati JS, Chronik BA, Wong D, Duggal N, et al. 1H MR spectroscopy of the motor cortex immediately following transcranial direct current stimulation at 7 Tesla. PLoS One. 2018;13(8):1–14.

60. Dwyer GE, Craven AR, Hirnstein M, Kompus K, Assmus J, Ersland L, et al. No Effects of Anodal tDCS on Local GABA and Glx Levels in the Left Posterior Superior Temporal Gyrus. Front Neurol. 2019;9(January):1–10.

61. Auvichayapat P, Aree-uea B, Auvichayapat N, Phuttharak W, Janyacharoen T, Tunkamnerdthai O, et al. Transient Changes in Brain Metabolites after Transcranial Direct Current Stimulation in Spastic Cerebral Palsy: A Pilot Study. Front Neurol. 2017;8(July):1–9.

62. Boissy P, Bourbonnais D, Kaegi C, Gravel D, Arsenault BA. Characterization of global synkineses during hand grip in hemiparetic patients. Arch Phys Med Rehabil. 1997;78(10):1117–24.

63. Nitsche MA, Paulus WJ. Excitability changes induced in the human motor cortex by weak transcranial direct current stimulation. J Physiol. 2000;527(3):633–9.

64. Antal A, Kincses TZ, Nitsche MA, Paulus WJ. Manipulation of phosphene thresholds by transcranial direct current stimulation in man. Exp Brain Res. 2003;150(3):375–8.

65. Boggio PS, Castro LO, Savagim EA, Braite R, Cruz VC, Rocha RR, et al. Enhancement of non-dominant hand motor function by anodal transcranial direct current stimulation. Neurosci Lett. 2006;404(1–2):232–6.

66. Anguera JA, Russell CA, Noll DC, Seidler RD. Neural correlates associated with intermanual transfer of sensorimotor adaptation. Brain Res. 2007;1185(1):136–51.

67. Lee M, Hinder MR, Gandevia SC, Carroll TJ. The ipsilateral motor cortex contributes to cross-limb transfer of performance gains after ballistic motor practice. J Physiol. 2010;588(1):201–12.

